# *IN VITRO* ANTI-VIRAL ACTIVITY OF HEXETIDINE (BACTIDOL^®^) ORAL MOUTHWASH AGAINST HUMAN CORONAVIRUS OC43 AND INFLUENZA A (H1N1) VIRUS

**DOI:** 10.1101/2021.02.11.430728

**Authors:** Marohren C Tobias -Altura, Corazon A Ngelangel

## Abstract

Mouthwashes are used to decrease oral cavity microbial load due to their antiseptic properties. Hexetidine is a broad-spectrum antiseptic used for minor infections of mucous membranes, and in particular as a 0.1% mouthwash for local infections and oral hygiene.

This study determined the anti-viral activity of the mouthwash hexetidine (Bactidol^®^), specifically in reducing viral concentration of Human Coronavirus OC43 (HCoV OC43; ATCC^®^ VR-1558™) and Influenza A virus (IAV H1N1; clinical strain) in Vero 6 and MDCK cell cultures respectively, using in-vitro suspension assay (ASTM E-1052-11) designed to evaluate virucidal property of microbicides like hexetidine.

Study results indicated that hexetidine was able to reduce infectivity of HCoV OC43 and IAV H1N1 at 25%, 50% and 100% concentrations by more than 80% at 15- and 30-seconds exposure times. One hundred percent (100%) concentration of hexetidine was found to be cytotoxic to MDCK cell line used for IAV H1N1 propagation. Hexetidine-treated cell lines achieved >80% survival rate for MDCK and Vero E6 at a contact time of 15 seconds and 30 seconds (which are the approximate times of gargling with hexetidine mouthwash).

The anti-viral activity of hexetidine mouthwash against other more virulent or pathogenic coronaviruses like SARS-CoV-2 can be explored further.

## INTRODUCTION

Mouthwashes have been widely used in oral hygiene. Most of the mouthwashes are used to decrease oral cavity microbial load due to their anti-septic properties and have been prescribed in dentistry. The active ingredients of mouthwashes, among others are hexetidine, benzydamine hydrochloride, and povidone iodine.

Hexetidine belongs to the group of pyrimidine derivatives. It is a broad-spectrum antiseptic, active *in vitro* and *in vivo* against Gram-positive and Gram-negative bacteria. It is used for minor infections of mucous membranes, and in particular as a 0.1% mouthwash for local infections and oral hygiene. Pharmacological evidence indicated that the primary mode of action is due to its interference with vital metabolic processes necessary for the growth of microorganisms [1].

Povidone iodine is an iodophore that is used as a disinfectant and antiseptic. Iodophores are loose complexes of iodine and carrier polymers. Solutions of povidone-iodine gradually release iodine to exert an effect against bacteria, fungi, viruses, protozoa, cysts, and spores. A 1% mouthwash has been used for oral infections including candidiasis [1].

Antimicrobial efficacy may be measured directly by the effect of certain compounds on the growth of microorganisms *in vitro*, or indirectly by studying their effect on certain health conditions that are caused or affected by microorganisms. Some of these conditions are halitosis, plaque, gingivitis and periodontitis. [1]

From the literature review above [2], hexetidine and povidone iodine generally had good efficacy against the bacterial flora of the oral cavity, with hexetidine being slightly superior between the two. In addition, hexetidine lost only approximately 25% of its antimicrobial action after 60 minutes while povidone iodine lost almost 40% of its antimicrobial efficacy after 10 minutes [3].

The sustained antimicrobial efficacy of hexetidine was also noted in another study [4]; however, it was noted that the duration was shorter than for chlorhexidine [3,4]. Hexetidine was also found to have good efficacy against plaque and gingivitis [5,6] but was again inferior to chlorhexidine [7].

Four other studies found that the antimicrobial action of povidone iodine had a short duration [3,11–13]. Some studies, however, found povidone iodine had higher antimicrobial efficacy than chlorhexidine [14,15]. Povidone iodine was also noted to reduce *Streptococcus mutans* count although not to the same degree as chlorhexidine [16]. It was more effective in alleviating mucositis than chlorhexidine [8] and was as effective as chlorhexidine for plaque and gingivitis [17].

The literature review above [2] concluded based on the *in vitro*, *in vivo* and clinical studies discussed, which used both direct and indirect methods to measure antimicrobial efficacy, that hexetidine, chlorhexidine and povidone-iodine were effective against oral microbial flora and differ primarily in the duration of their antimicrobial action.

Zoonotic coronaviruses were discovered in the 1960s and pathogenic human coronaviruses were discovered in 2002. Currently, there are now seven human coronaviruses which include the SARS-CoV-2. Most of these human coronaviruses cause mild diseases, and these include the HCoVs OC43, 229E, NL63 and HKU1. The more virulent species are SARS-CoV, MERS-CoV and SARS-CoV-2. Coronaviruses are positive sense, single stranded RNA virus, spherical and enveloped with club-shaped spikes on the surface looking like the solar corona. The four coronaviruses genera are α, β, γ, δ. The human α-CoV are HCoV-229E and NL63, while the human β-CoV are MERS-CoV, SARS-CoV, HCoV OC43 and HCoV-HKU1. SARS-CoV-2 and HCoV OC43 are both β-CoV.[18]

This study determined the anti-viral activity of hexetidine (Bactidol^®^), specifically in reducing viral concentration of Human Coronavirus (HCoV OC43) and Influenza A virus (IAV H1N1) infected cell cultures by demonstrating its Tissue Culture Infective Dose or TCID50/ml in 25%, 50% and 100% aqueous hexetidine concentration at 15- and 30-seconds exposure times (which are the approximate times of gargling with Bactidol^®^ mouthwash).

## MATERIALS and METHODS

The study was done from 10 September 2020 to 03 November 2020. The test method used was according to the American Society for Testing and Materials International - ASTM E-1052-11 Standard test method, which assess the activity of microbicides against viruses in suspension (2011) [19].

### Tissue culture cells

Madin Darby Canine Kidney (MDCK) cells from WHO Collaborating Centre for Reference and Research on Influenza, Australia, and African green monkey kidney Vero E6 cells from the Department of Virology, Tohoku University Graduate School of Medicine, Sendai, Japan were used. These cell lines are permissive and susceptible for the test viruses IAV H1N1 and HCoV OC43, respectively. These cultures were maintained in tissue culture media with additives.

### Test viruses

Influenza A virus H1N1 (IAV H1N1), came from a clinical sample from Influenza Surveillance, were grown in MDCK cells, confirmed by Hemagglutination Inhibition and RT-PCR. Human Coronavirus (HCoV FR-302), Strain OC43 (ATCC^®^VR-1558™) was purchased from American Type Culture Collection (ATCC). These are the common respiratory viruses that can cause sore throat and are found in the oral cavity; HCoV is a β-CoV similar to SARS-CoV-2. These viruses were grown on viral culture media with additives (L-glutamine, Penicillin-Streptomycin).

### Antiviral agents

Hexetidine (Bactidol^®^) at 100% (undiluted), aqueous 50% and 25% concentrations were used. 70% ethyl alcohol served as a positive control.

The experimental conditions were: test temperature at 37°C; neutralizer used, Minimum Essential Medium (Gibco brand, Catalogue No. 11700-077) with 2% Fetal Bovine Serum (heat inactivated, Gibco Brand, Catalogue No. 10500-064); incubation time for IAV H1N1 - 5 days, HCoV OC43 - 3 days; and incubation temperature for IAV H1N1 - 35°C, HCoV OC43 – 37°C.

The method used was an *in-vitro* suspension assay designed to evaluate virucidal property of the product against IAV H1N1 and HCoV OC43. Presence of infective virus was determined via monitoring of cytopathic effect (CPE) on the appropriate cell line. A 0.2 ml suspension of virus was exposed to 1.8mI per specified concentration of anti-viral test agents. At each specified exposure time, an aliquot was removed and neutralized by serial dilution and assayed for presence of virus. Virus controls, cytotoxicity controls, and neutralization controls were assayed in parallel.

After incubation, virus infected cell cultures in wells were identified through their characteristic CPE, rounding of cells (MDCK) and vacuolization of the cytoplasm and sloughing of cells (Vero E6). Recorded results were used to calculate infecting activity (TCID50) through Spearman-Karber method [20]. Percent reduction and log reduction were subsequently computed.

All tests were done at the Virology Department of the Research Institute for Tropical Medicine (RITM), Department of Health. The Institutional Review Board (IRB) of RITM was accordingly informed of this basic laboratory study involving viral cell cultures and does not require to undergo review as per regulations on studies involving animals or humans as test subjects.

## RESULTS

The results of the ASTM E-1052-11 assay, hexetidine (Bactidol^®^) against IAV H1N1 (clinical strain) and HCoV OC43 (ATCC^®^ VR-1558™) are shown in Table 1. Hexetidine (Bactidol^®^) at 25%, 50% and 100% aqueous concentrations against HCoV OC43 (ATCC^®^ VR-1558™) showed from 94.38 % to 99.68% infectivity reduction at 15 seconds contact time, and from 94.38% to 99% infectivity reduction at 30 seconds contact time.

**Table 1.**
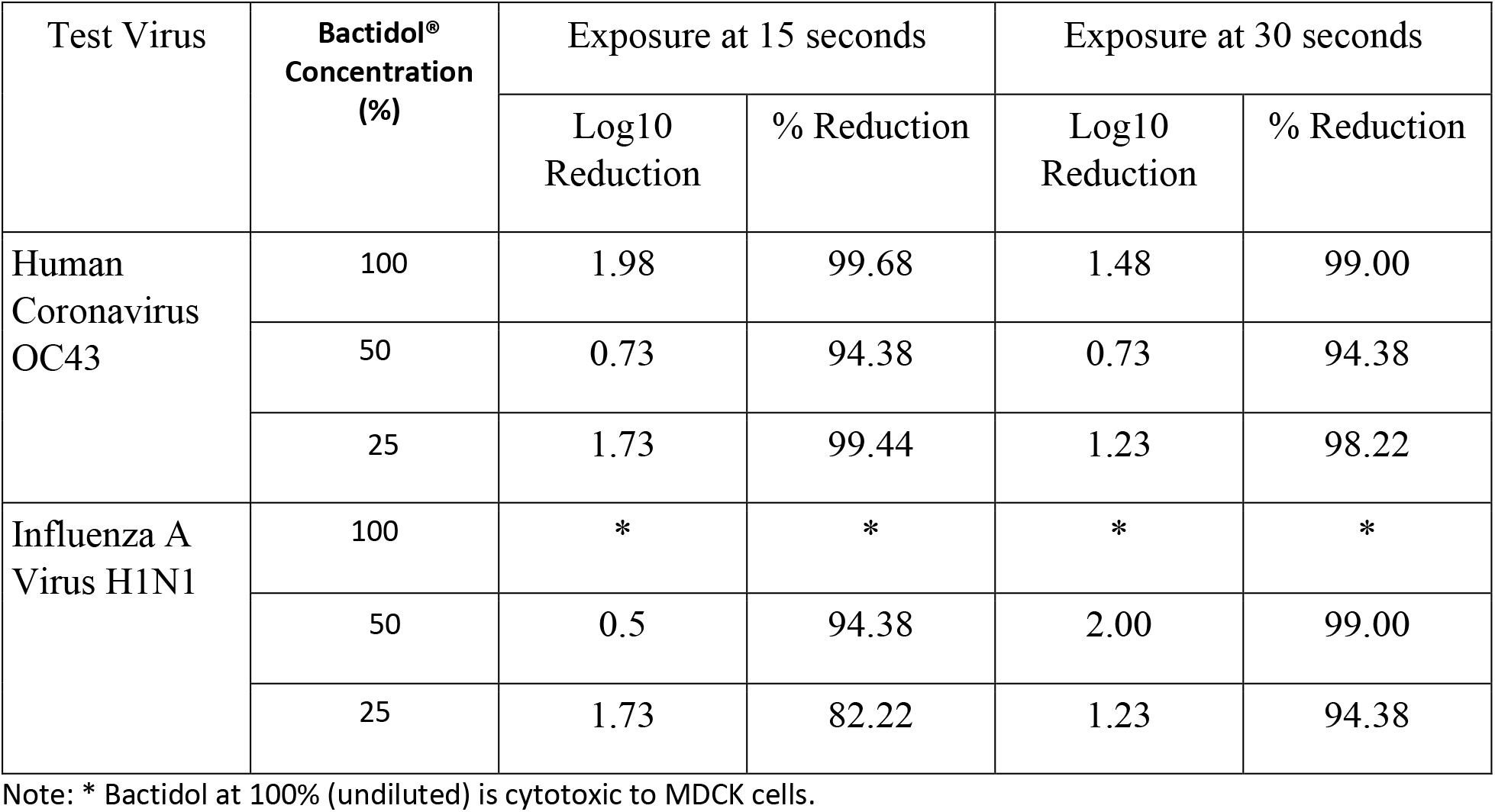
Computed Log10 reduction and percent (%) reduction values for IAV H1N1 (clinical strain) and HCoV OC43 (ATCC^®^ VR-1558™) challenged with three concentrations of hexetidine (Bactidol^®^) at 15- and 30-seconds exposure times.

| Test Virus | Bactidol® Concentration (%) | Exposure at 15 seconds |  | Exposure at 30 seconds |  |
| --- | --- | --- | --- | --- | --- |
|  |  | Log10 Reduction | % Reduction | Log10 Reduction | % Reduction |
| Human Coronavirus OC43 | 100 | 1.98 | 99.68 | 1.48 | 99.00 |
|  | 50 | 0.73 | 94.38 | 0.73 | 94.38 |
|  | 25 | 1.73 | 99.44 | 1.23 | 98.22 |
| Influenza A Virus H1N1 | 100 | * | * | * | * |
|  | 50 | 0.5 | 94.38 | 2.00 | 99.00 |
|  | 25 | 1.73 | 82.22 | 1.23 | 94.38 |

Note: \* Bactidol at 100% (undiluted) is cytotoxic to MDCK cells.

Hexetidine (Bactidol^®^) at 25% and 50% aqueous concentrations against IAV H1N1 (clinical strain) showed 82.22% and 94.38% infectivity reduction at 15 seconds contact time, and from 94.38% and 99% infectivity reduction at 30 seconds contact time, at respective concentrations.

However, one hundred percent (100%) concentration of hexetidine (Bactidol^®^) was found to be cytotoxic to the MDCK cell line used for IAV H1N1 propagation.

The following figures were taken during the study, Hexetidine (Bactidol^®^) mouthwash showed inhibition of the virus induced CPE in the infected MDCK and Vero 6 cells, Figure 1. letters A and C, respectively.

**Figure 1.**
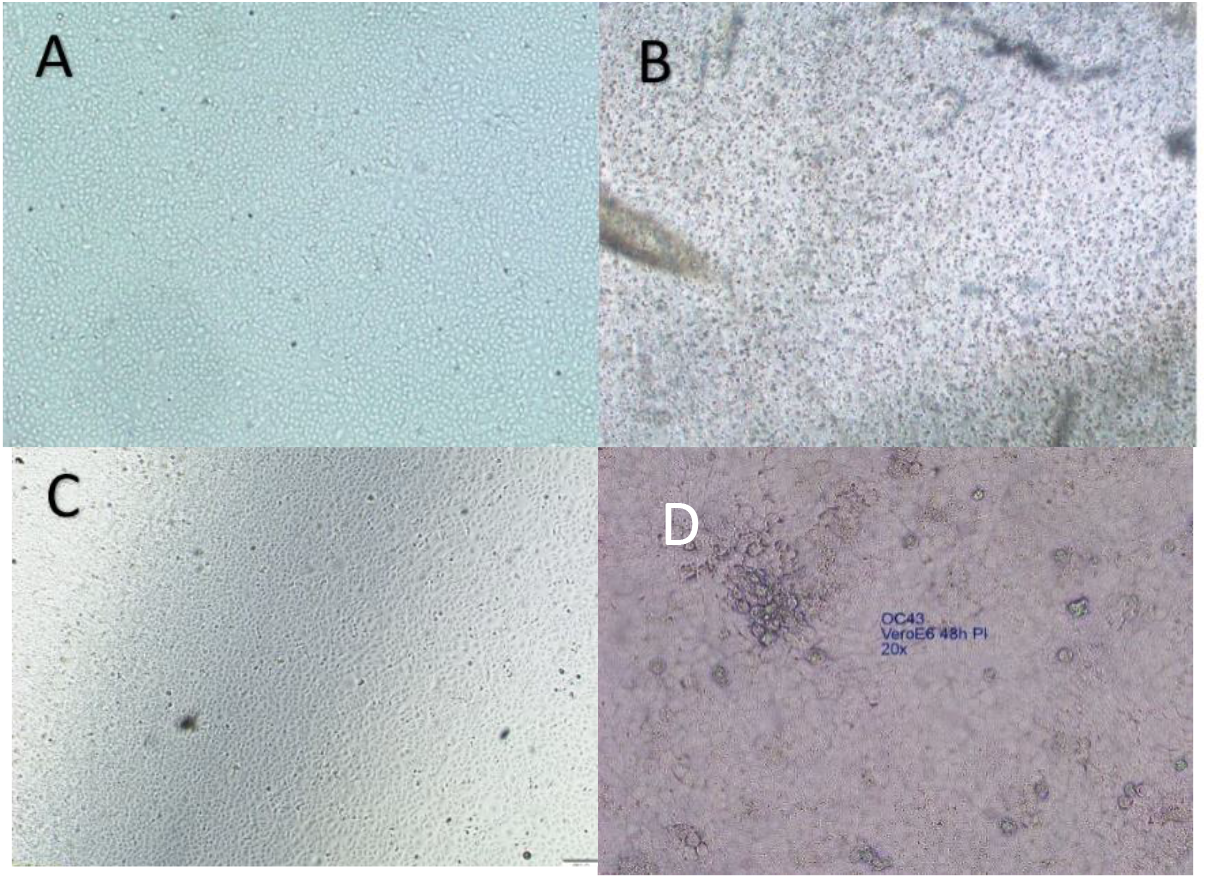
**(A)** confluent MDCK monolayer without IAV CPE, **(B)** MDCK monolayer infected with IAV exhibited rounding of cells cytopathic effect; **(C)** confluent Vero E6 monolayer without HCoV OC43 CPE; **(D)** Vero E6 monolayer infected with HCoV OC43 exhibited vacuolization of the cytoplasm and sloughing of cells cytopathic effect

## DISCUSSION

This study showed that hexetidine (Bactidol^®^) was able to reduce infectivity of HCoV OC43 (ATCC^®^ VR-1558™) and IAV H1N1 at 25%, 50% and 100% concentrations. These hexetidine concentrations reduced by more than 94% to 99% the infectivity of Human Coronavirus OC43 at 15 and 30 seconds exposure times.

For Influenza A Virus H1N1 (clinical strain) showed that at lower, 25% hexetidine concentration 82.22% and 94.38% infectivity reduction was found at 15 and 30 seconds contact time, respectively. At 50% concentration, 94.38% and 99% infectivity reduction at 15 and 30 seconds contact time, respectively. Undiluted hexetidine (Bactidol^®^), 100% concentration was cytotoxic to the MDCK cell line used for Influenza A H1N1 propagation.

Hexetidine mouthwash inhibited influenza A virus (IAV) and human coronavirus (HCoV OC43). Thus, hexetidine (Bactidol^®^) in 25%, 50%, and 100% concentrations can be used as an effective mouthwash over 15 to 30 seconds to get rid of HCoV OC43 and IAV H1N1 on the oral mucosal surfaces.

The study of Deryabin P.G., et al., “Analysis of Antiviral Properties of Hexoral *In Vitro* against Some Viruses that Cause Acute Respiratory Infections and Herpes” has shown similar results. Hexoral^®^ (with Hexetidine as the active ingredient) has shown to have an antiviral property against viruses causing human respiratory tract infections and herpes virus. [21] Exposure to Hexoral^®^ and Hexetidine alone for 30 secs. was able to attenuate the infectivity of Influenza Virus A/H5N1, pandemic Influenza Virus A/H1N1pdm, Respiratory Syncytial Virus, and Herpes Simplex Virus type 1 by 100 or more times. [21]

Antiseptic mouthwashes have been widely used as a standard measure before routine dental treatment, especially pre-operatively [22,23]. These are widely used solutions for rinsing the mouth due to their ability to reduce the number of microorganisms in the oral cavity and colony-forming units in dental aerosols [24]. The use of mouthwashes and gargles were deemed to be relatively safe, with only minor adverse effects with long-term use [25].

Vergara-Buenaventura A and Castro-Ruiz C (2020) suggested the use of pre-procedural mouthwashes in dental practice to reduce SARS-CoV-2 viral load from dental procedures and to reduce the cross-infection risk while treating patients during the pandemic (26).

The SARS-CoV-2 spike protein in the membrane envelope is a typical structure of coronaviruses [26–28]. The spike protein interacts with the angiotensin-converting enzyme 2 (ACE2) receptors of the host cells enabling the virus to enter the cells [29]. Different tissue cells, including those of the mucosal tissues, gingiva, periodontal pockets, tongue and salivary glands present possible infection routes as these have membranes bound to ACE2 [29–34]. Such oral tissues are sources from which SARS-CoV-2 transmission may occur during dental care, talking, coughing, and sneezing (35,36).

The American Dental Association [37] and the Center for Disease Control and Prevention [38] have recommended the use of pre-procedural mouthwashes before oral procedures, despite no clinical evidence that the use of mouthwashes could prevent SARS-CoV-2 transmission. [26]

In the Philippines, among the active agents used in commercially available mouthwashes, only povidone-iodine had been shown to have antiviral activity *in vitro* against SARSCoV-2 [39].

This *in vitro* study showed that hexetidine, even at diluted concentrations reduced the infectivity of the two oral virus strains, HCoV OC43 and Influenza A virus H1N1 when used for 15 and 30 seconds. The anti-viral activity of hexetidine mouthwash against the other more virulent members of the Coronavirus Family, SARS-CoV-2 can be explored using the methods used in this *in vitro* study. Clinical studies can also follow, to evaluate the effectiveness of antiseptic mouthwashes like hexetidine on SARS-CoV-2, the subjects may be with asymptomatic or mildly infected individuals. The future study will determine whether hexetidine (Bactidol^®^) can lessen viral load or shorten the duration of viral carriage.

## ACKNOWLEDGEMENT

Thanks is due to the Research Institute for Tropical Medicine-Department of Health and its Virology Department Staff Mayan U Lumandas, Vina Lea F Arguelles, Ma Therese DC Quimpo, and Janiza Ianne M Foronda, who did the laboratory tests of this study.

## DISCLAIMER

This in-vitro study initiative has been funded by Johnson and Johnson (Philippines), Incorporated through a research grant to Asian Hospital, Inc.

## References

1. Sweetman S. Martindale, the Complete Drug Reference. 36th edition, Grayslake, IL; 2009.

2. Maldo D, Tomas A (Medical Affairs-iMEDGlobal-Philippines). Hexetidine, Benzydime, Povidone Iodine on Oral Germ Kill Literature Search & Review for Johnson and Johnson Philippines-Medical Affairs, with literature search from 21-22 May 2015 (yielding 19 articles from Ovid and PubMed). 2015. (J&J Data on File)

3. Pitten F A, Kramer A. Antimicrobial efficacy of antiseptic mouth rinse solutions. Eur J Clin Pharmacol. 1999;55(2):95–100.

4. Polat H.B., Ozdemir H., Ay S. Effect of different mouth rinses on third molar surgery-related oral malodor. Oral Surgery, Oral Medicine, Oral Pathology, Oral Radiology and Endodontology. 105 (3) (pp e1–e8), 2008. Date of Publication: March 2008.

5. Ernst C.-P., Canbek K., Dillenburger A., Willershausen B. Clinical study on the effectiveness and side effects of hexetidine and chlorhexidine mouth rinses versus a negative control. Quintessence Int. 2005;36:641–652.

6. Sharma N.C., Galustians H.J., Qaqish J., et al. Antiplaque and anti-gingivitis effectiveness of a hexetidine mouthwash. Journal of clinical periodontology. 2003;30 (7):590–594.

7. Afennich F, Slot D E, Hossainian N, Van der Weijden G.A. The effect of hexetidine mouthwash on the prevention of plaque and gingival inflammation: a systematic review. International journal of dental hygiene. 2011;9(3):182–190.

8. Roberts WR; Addy M. Comparison of the *in vivo* and *in vitro* antibacterial properties of antiseptic mouth rinses containing chlorhexidine, alexidine, cetyl pyridinium chloride and hexetidine. Relevance to mode of action. Journal of Clinical Periodontology. 1981 Aug;8(4):295–310.

9. Samaranayake L.P., Robertson A.G., MacFarlane T.W., et al. The effect of chlorhexidine and benzydamine mouthwashes on mucositis induced by therapeutic irradiation. Clinical Radiology. 1988;39 (3):291–294.

10. Matthews RW; Scully CM; Levers BG; Hislop WS. Clinical evaluation of benzydamine, chlorhexidine, and placebo mouthwashes in the management of recurrent aphthous stomatitis. Oral Surgery, Oral Medicine, Oral Pathology. 1987 Feb; 63(2):189–91.

11. Altonen M, Saxen L, Kosunen T, Ainamo J. Effect of two antimicrobial rinses and oral prophylaxis on preoperative degerming of saliva. Int J Oral Surg. 1976;5(6):276–84.

12. Kosutic D., Uglesic V., Perkovic D., et al. Preoperative antiseptics in clean/contaminated maxillofacial and oral surgery: prospective randomized study. International Journal of Oral and Maxillofacial Surgery. 2009;38 (2):160–165.

13. Addy M; Wright R. Comparison of the *in vivo* and *in vitro* antibacterial properties of povidone iodine and chlorhexidine gluconate mouth rinses. Journal of Clinical Periodontology.1978 Aug; 5(3):198–205.

14. Yoneyama A, Shimizu M, Tabata M, Yashiro J, Takata T, Hikida M. *In vitro* short-time killing activity of povidone-iodine (Isodine Gargle) in the presence of oral organic matter. Dermatology (Basel). 2006;212 Suppl 1:103–8.

15. Shiraishi T, Nakagawa Y. Evaluation of the bactericidal activity of povidone-iodine and commercially available gargle preparations. Dermatology (Basel). 2002;204 Suppl 1:37–41.

16. Neeraja R., Anantharaj A., Praveen P, et al. The effect of povidone-iodine and chlorhexidine mouth rinses on plaque Streptococcus mutans count in 6- to 12-year-old school children: An *in vivo* study. Journal of Indian Society of Pedodontics and Preventive Dentistry.2008; 26 (SUPPL. 5):S14–S18.

17. Fine PD. A clinical trial to compare the effect of two antiseptic mouthwashes on gingival inflammation. Fine PD. Journal of Hospital Infection. 1985 Mar;6 Suppl A:189–93.

18. Malik YA. Properties of Coronavirus and SARS-CoV-2. Malaysian J Pathol. 2020;42(1): 3–11

19. American Society for Testing and Materials. ASTM E-1052-11 Standard test method to assess the activity of microbicides against viruses in suspension (2011). (http://www.materialstandard.com/wp-content/uploads/2019/10/E1052-11.pdf accessed 17/12/2020).

20. Hamilton MA, Russo RC, Thurston RV. Trimmed. Spearman-Kärber CPE Method for Estimating Median Lethal Concentrations in Toxicity Bioassays. Environ Sci Technol. 1977;11:714–719. doi

21. Deryabin PG, Galegov GA, Andronova VA, and Botikov AG. Analysis of Antiviral Properties of Hexoral *In Vitro* against Some Viruses that Cause Acute Respiratory Infections and Herpes. Bulletin of Experimental Biology and Medicine. 2016 Jan; 160 (3) (Translated from Byulleten’ Eksperimental’noi Biologii i Meditsiny. 2015 Sept; 160 (9): 339–342).

22. Kosutic D, Uglesic V, Perkovic D, et al. Preoperative antiseptics in clean/contaminated maxillofacial and oral surgery: prospective randomized study. Int J Oral Maxillofac Surg. 2009;38:160–5.

23. Dominiak M, Shuleva S, Silvestros S, et al. A prospective observational study on perioperative use of antibacterial agents in implant surgery. AdvClin Exp Med. 2020;29:355–63.

24. Marui VC, Souto MLS, Rovai ES, et al. Efficacy of preprocedural mouth rinses in the reduction of microorganisms in aerosol: a systematic review. J Am Dent Assoc. 2019;150:1015–26, e1.

25. DePaola L, Spolarich E. Safety and efficacy of antimicrobial mouth rinses in clinical practice. Access. 2007;81(5):13–25.

26. Vergara-Buenaventura A, Castro-Ruiz C. Use of mouthwashes against Covid-19 in dentistry. British J of Oral & Maxillofacial Surg. 2020;58: 924–927 (with cross-references).

27. Yoon JG, Yoon J, Song JY, et al. Clinical significance of a high SARS-CoV-2 viral load in the saliva. J Korean Med Sci. 2020;35:e195.

28. Li F. Structure, function, and evolution of coronavirus spike proteins. Annu Rev Virol. 2016;3:237–61.

29. Chen Y, Guo Y, Pan Y, et al. Structure analysis of the receptor binding of 2019-nCoV. Biochem Biophys Res Commun. 2020 Feb;525:135–40.

30. Xu H, Zhong L, Deng J, et al. High expression of ACE2 receptor of 2019-nCoV on the epithelial cells of oral mucosa. Int J Oral Sci. 2020;12:8.

31. Wan Y, Shang J, Graham R, et al. Receptor recognition by the novel coronavirus from Wuhan: an analysis based on decade-long structural studies of SARS coronavirus. J Virol. 2020;94:e00127–220.

32. Hamming I, Timens W, Bulthuis ML, et al. Tissue distribution of ACE2 protein, the functional receptor for SARS coronavirus. A first step in understanding SARS pathogenesis. J Pathol. 2004;203:631–7.

33. Li Y, Ren B, Peng X, et al. Saliva is a non-negligible factor in the spread of COVID-19. Mol Oral Microbiol. 2020;35:141–5.

34. Badran Z, Gaudin A, Struillou X, et al. Periodontal pockets: a potential reservoir for SARS-CoV-2? Med Hypoth. 2020;143:109–907.

35. Siqueira WL, Moffa EB, Mussi MC, et al. Zika virus infection spread through saliva – a truth or myth? Braz Oral Res. 2016;30. S1806–83242016000100801.

36. Anschau V, Sanjuán R. Fibrinogen gamma chain promotes aggregation of vesicular stomatitis virus in saliva. Viruses. 2020;12:282.

37. American Dental Association. ADA interim guidance for minimizing risk of COVID-19 transmission; 2020. Available from URL:https://www.kavo.com/en-us/resource-center/ada-interim-guidance-minimizing-risk-covid-19-transmission [accessed 13.08.20].

38. Centers for Disease Control and Prevention. Interim infection prevention and control guidance for dental settings during the COVID-19 response. Available from URL: https://www.cdc.gov/coronavirus/2019-ncov/hcp/dental-settings.html [accessed 13.08.20].

39. Bidra AS, Pelletier JS, Westover JB, Frank S, Brown SM, Tessema B. Rapid In‐Vitro Inactivation of Severe Acute Respiratory Syndrome Coronavirus 2 (SARS‐CoV‐2) Using Povidone‐Iodine Oral Antiseptic Rinse. J Prosthodont. 2020;2:1–14.

